# A role for tetracycline selection in the evolution of *Clostridium difficile* PCR-ribotype 078

**DOI:** 10.1101/262352

**Authors:** Kate E. Dingle, Xavier Didelot, T. Phuong Quan, David W. Eyre, Nicole Stoesser, Charis A. Marwick, John Coia, Derek Brown, Sarah Buchanan, Umer Z. Ijaz, Cosmika Goswami, Gill Douce, Warren N. Fawley, Mark H. Wilcox, Timothy E.A. Peto, A. Sarah Walker, Derrick W. Crook

## Abstract

Farm animals have been identified as reservoirs of *Clostridium difficile* PCR-ribotype 078 (RT078). Since 2005, the incidence of human clinical cases (frequently severe), with this genotype has increased. We aimed to understand this change, by studying the recent evolutionary history of RT078. Phylogenetic analysis of international genomes (isolates from 2006–2014) revealed several recent clonal expansions. A common ancestor of each expansion had independently acquired different alleles of the tetracycline resistance gene *tetM*. Consequently, an unusually high proportion of RT078 genomes were *tetM* positive (76.5%). Additional tetracycline resistance determinants were also identified, some for the first time in *C. difficile* (efflux pump *tet40*). Each *tetM*-clonal expansion lacked geographic structure, indicating rapid international spread. Resistance determinants for *C. difficile*-infection-triggering antimicrobials including fluoroquinolones and clindamycin were comparatively rare in RT078. Tetracyclines are used intensively in agriculture; this selective pressure, plus rapid spread via the food-chain may explain the increased RT078 prevalence in humans.

## Introduction

*Clostridium difficile* infection is a significant international challenge, affecting patients in community and healthcare environments worldwide (Davies et al., 2014; Lessa et al., 2015; Wilcox et al., 2008). The severity of symptoms ranges from mild diarrhoea to pseudomembranous colitis and toxic megacolon. Crude 30-day mortality in the UK is 16% (in an endemic setting) and can exceed 30% (Planche et al., 2013; McGowan et al., 2011), while in the US it has been estimated that almost half a million CDIs caused 29,000 deaths in a single year (Lessa et al., 2015).

The molecular epidemiology of *Clostridium difficile* infection varies both temporally and geographically, frequently in response to local antimicrobial prescribing (Lessa et al., 2015; Public Health England, 2016; Wilcox et al., 2012; Dingle et al., 2017). Clinically important outbreak-associated genotypes can emerge when the inherent resistance of *C. difficile* to cephalosporins (Shuttleworth et al., 1980) is supplemented with acquired resistance to certain high risk antimicrobials, including clindamycin (Johnson et al., 1999) and more recently fluoroquinolones. The latter contributed to the emergence of multiple phylogenetically unrelated outbreak-associated genotypes including the ‘hypervirulent’ PCR-ribotype 027 (Loo et al., 2005; He et al., 2017; Spigaglia., 2016; Dingle et al., 2017). However, the reason(s) for the changing prevalence of other clinically important *C. difficile* genotypes is frequently unknown (Belmares et al., 2009). These include RT078, which is unusual in having an agricultural link (Goorhuis et al., 2008).

The increased importance of *C. difficile* RT078 as a human pathogen was first reported in The Netherlands, rising from 3% to 13% of *Clostridium difficile* infection cases during 2005–2008 (Goorhuis et al., 2008). Around the same time, a ten fold increase was noted in North America (Jhung et al., 2008). Similar increases and occasional outbreaks were subsequently recorded throughout Europe (Barbut et al., 2007; Bauer et al., 2011; Burns et al., 2010) and *C. difficile* RT078 has recently increased to 4.4%, 9.7% and 8.1% of total *Clostridium difficile* infection cases in North America, England and Scotland, respectively (Lessa et al., 2015; Mulvey et al., 2010; Fawley et al., 2016; Health Protection Scotland (2017). Three distinctive features of RT078-associated *Clostridium difficile* infection raise specific concerns, namely: increased severity of disease with the highest genotype-specific mortality rate (Goorhuis et al., 2008; Walker et al., 2013), a higher proportion of community-associated disease, and more infections in younger age groups compared with other genotypes (Lessa et al., 2015; He et al., 2013; Taori et al., 2014).

The agricultural association of *C. difficile* RT078 reflects its isolation from sick and healthy animals (frequently pigs), bird droppings, vermin and the farm environment (Hensgens et al., 2012; Keel et al., 2007; Bandelj et al., 2014; Burt et al., 2012). However, in common with many other toxin producing *C. difficile* genotypes, ribotype RT078 can be carried asymptomatically by human infants and adults (Stoesser et al., 2017; Knetsch et al., 2014). This genotype has also been isolated from a variety of retail meat products including pork, beef, and others (Curry et al., 2012; Songer et al., 2007). Therefore the natural reservoirs of RT078 support the hypothesis that humans become colonised via the food chain and/or the environment (Hensgens et al., 2012).

Whole genome sequence data have been used to study the emergence and transmission of many bacterial pathogens. The international dissemination of ‘hypervirulent’ fluoroquinolone resistant *C. difficile* 027 was revealed in this way (He et al., 2013) and its rapid localised nosocomial transmission was demonstrated, as for other fluoroquinolone resistant genotypes (Dingle et al., 2017). Here, we used whole genome sequencing and phylogenetic approaches to study the recent evolutionary history of *C. difficile* RT078 and investigate the hypothesis that the recent clinical prominence of this genotype is due to antimicrobial selection.

## Results

The role of antimicrobial selection in the emergence of *C. difficile* RT078 was investigated using a large, international collection of genomes from the UK, Europe and North America (Table S1, n=400, Dingle et al., 2017; Stoesser et al., 2017; Knetsch et al., 2014; Louie et al., 2011; Cornely et al., 2012). Virtually all RT078 *C. difficile* share the same multilocus sequence type, ST11 (Dingle et al., 2011). However, this ST also includes the very closely related PCR-ribotypes RT126, RT033, RT045 and RT066 (relationship to RT078 shown phylogenetically, Figure S1). These ST11-associated PCR-ribotypes (RT126, RT033, RT045 and RT066) are relatively rare clinically. Only one RT078 genome included was a single locus variant of ST11; ST317. The overall proportion of RT078 within the collections from which the study genomes were sourced (see Materials and Methods) was 3–4%, because these data sets are dominated by RT027. Genomes studied are referred to subsequently as RT078. The presence of determinants of resistance to tetracyclines (*tetM*), fluoroquinolones (*gyrA*/*B* substitutions), aminoglycosides (*aphA1* or AAC(6’)-APH(2’)) and clindamycin (*ermB*) within these genomes was assessed (Table 1).

**Table 1.**
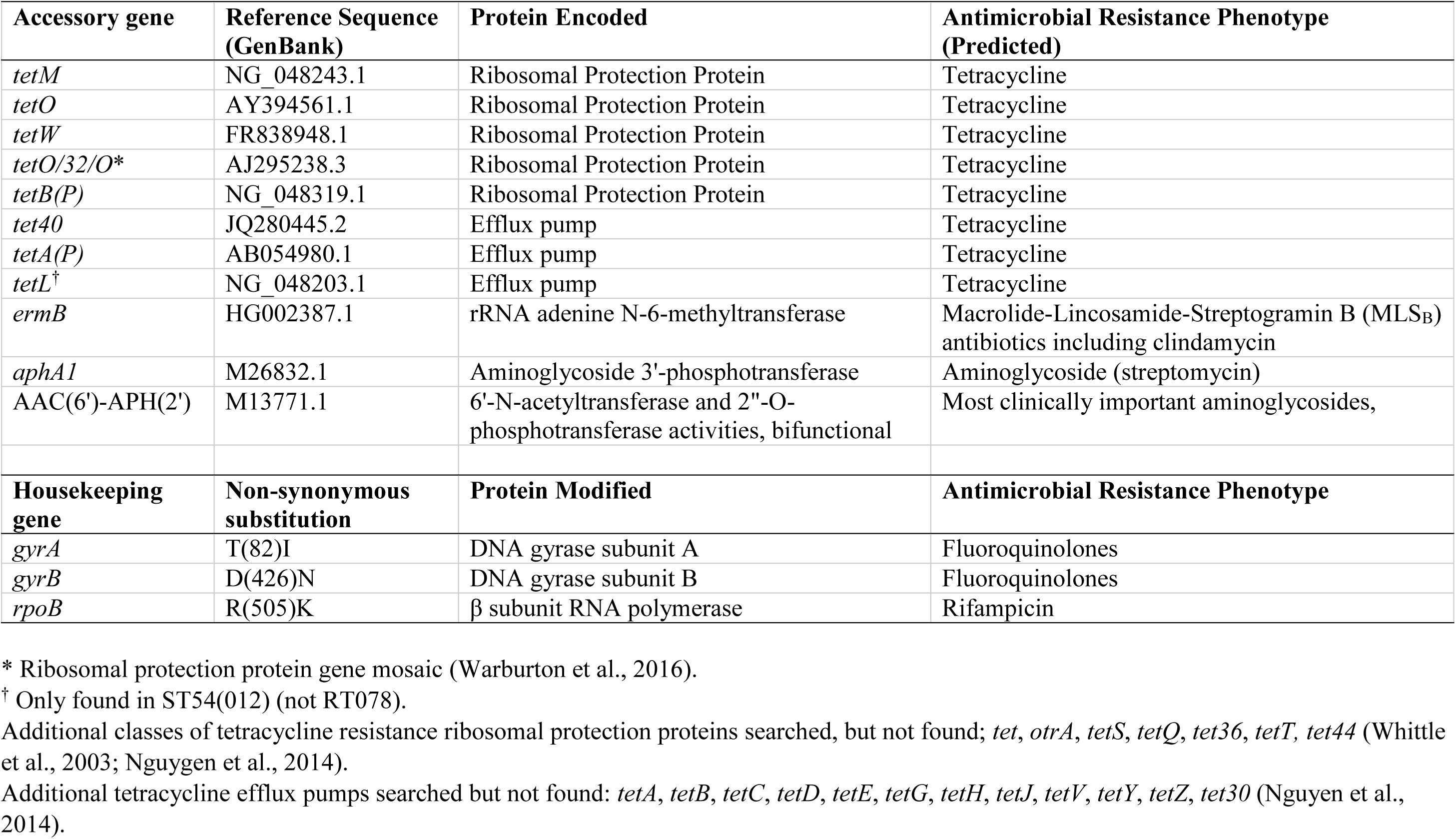
Antimicrobial resistance genes used to search *C. difficile* whole genome sequences

| Accessory gene | Reference Sequence (GenBank) | Protein Encoded | Antimicrobial Resistance Phenotype (Predicted) |
| --- | --- | --- | --- |
| *tetM* | NG_048243.1 | Ribosomal Protection Protein | Tetracycline |
| *tetO* | AY394561.1 | Ribosomal Protection Protein | Tetracycline |
| *tetW* | FR838948.1 | Ribosomal Protection Protein | Tetracycline |
| *tetO/32/O\** | AJ295238.3 | Ribosomal Protection Protein | Tetracycline |
| *tetB(P)* | NG_048319.1 | Ribosomal Protection Protein | Tetracycline |
| *tet40* | JQ280445.2 | Efflux pump | Tetracycline |
| *tetA(P)* | AB054980.1 | Efflux pump | Tetracycline |
| *tetL*<sup>†</sup> | NG_048203.1 | Efflux pump | Tetracycline |
| *ermB* | HG002387.1 | rRNA adenine N-6-methyltransferase | Macrolide-Lincosamide-Streptogramin B (MLS<sub>B</sub>) antibiotics including clindamycin |
| *aphA1* | M26832.1 | Aminoglycoside 3'-phosphotransferase | Aminoglycoside (streptomycin) |
| AAC(6')-APH(2') | M13771.1 | 6'-N-acetyltransferase and 2"-O-phosphotransferase activities, bifunctional | Most clinically important aminoglycosides, |
| Housekeeping gene | Non-synonymous substitution | Protein Modified | Antimicrobial Resistance Phenotype |
| *gyrA* | T(82)I | DNA gyrase subunit A | Fluoroquinolones |
| *gyrB* | D(426)N | DNA gyrase subunit B | Fluoroquinolones |
| *rpoB* | R(505)K | β subunit RNA polymerase | Rifampicin |

\* Ribosomal protection protein gene mosaic (Warburton et al., 2016).
<sup>†</sup> Only found in ST54(012) (not RT078).
Additional classes of tetracycline resistance ribosomal protection proteins searched, but not found; *tet*, *otrA*, *tetS*, *tetQ*, *tet36*, *tetT*, *tet44* (Whittle et al., 2003; Nguyen et al., 2014).
Additional tetracycline efflux pumps searched but not found: *tetA*, *tetB*, *tetC*, *tetD*, *tetE*, *tetG*, *tetH*, *tetJ*, *tetV*, *tetY*, *tetZ*, *tet30* (Nguyen et al., 2014).

### Prevalence of antimicrobial resistance determinants in *C. difficile* RT078

*tetM*, the presence of which leads to tetracycline resistant phenotype (Knetsch et al., 2014), was by far the most prevalent antimicrobial resistance determinant in RT078, at 77.5% (148/191) Oxfordshire and Leeds, 76.4% (84/110) Scottish and 75.0% (24/32) North American and European genomes (Table S1). *gyrA*/*B* substitutions, which reduce susceptibility to fluoroquinolones, were less prevalent at 13.0% (25/191) among Oxfordshire and Leeds RT078 genomes, 10.0% (11/110) among Scottish and 18.9% (6/32) in North American and European RT078s. Similarly, *aphA1* (aminoglycoside resistance) was detected in 21.5% (41/191) Oxfordshire and Leeds, 9.1% (10/110) Scottish and 40.6% (13/32) North American and European genomes. Finally, *ermB* (clindamycin resistance) occurred in 4.2% (8/191) Oxfordshire and Leeds, 4.5% (5/110) Scottish and 18.8% (6/32) North American and European RT078.

### Prevalence of *tetM* in the clinical *C. difficile* population

To determine whether the high *tetM* prevalence in RT078 was unusual among clinical *C. difficile*, all available additional genomes of other genotypes (defined by ST/PCR-ribotype), from the same isolate collections (except those comprising only RT078 – Scottish and Dutch (unpublished and Knetsch et al., 2014)), were examined for the presence of *tetM* (Figure 1A). The non-RT078 genotypes were described using the notation ST37(017) to indicate, for example, Multilocus Sequence Type 37 and the equivalent (PCR-ribotype 017).

**Figure 1.**
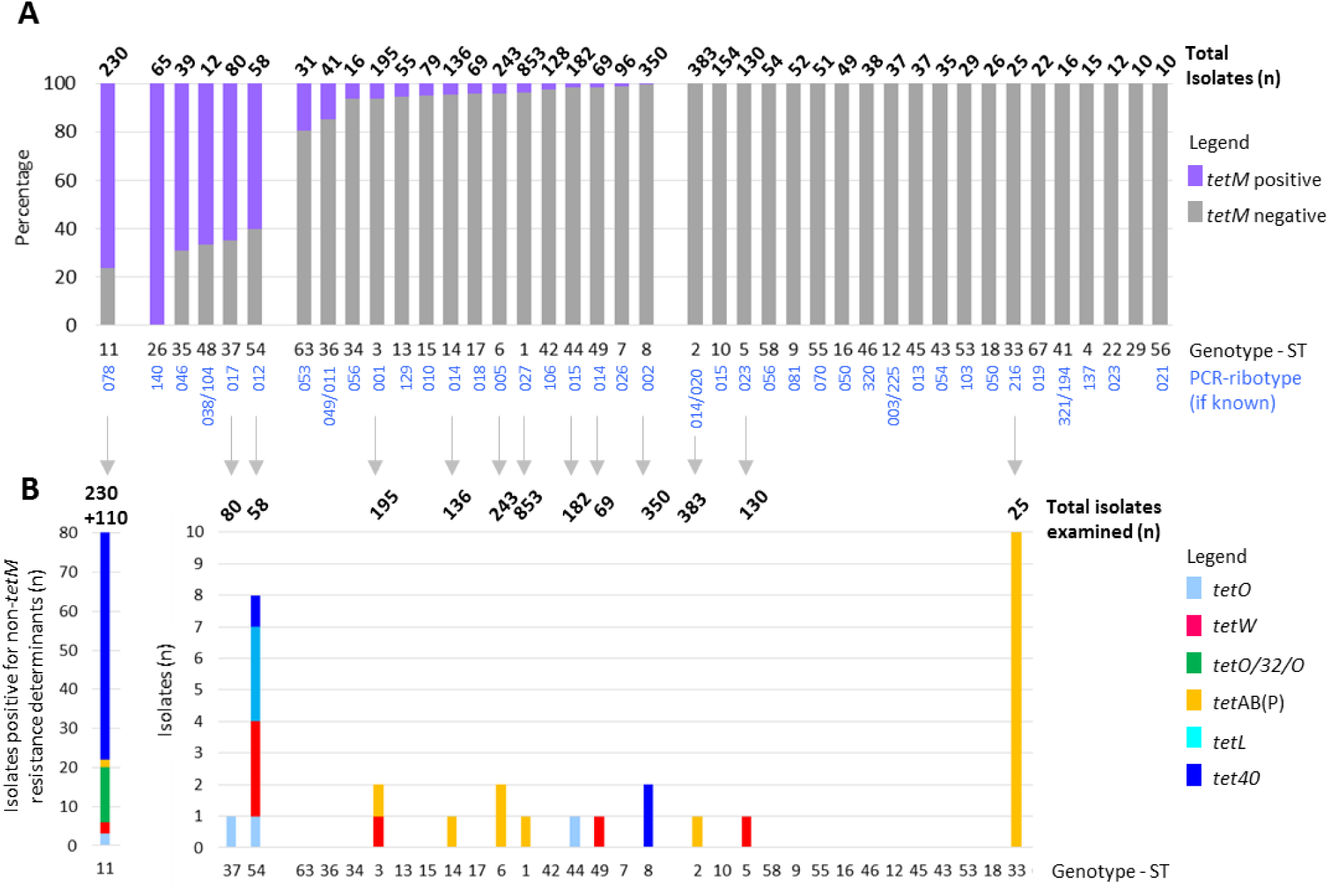
Prevalence of tetracycline resistance determinants in RT078 and other clinically relevant *C. difficile* genotypes. **(A) The proportion (%) of each clinically important genotype that was positive for the ribosomal protection protein (RPP) gene *tetM*.** Data are shown for genotypes having 10 genomes or more, from isolate collections representing Oxfordshire (EIA positives, negatives, infant and farm), Leeds, North America and Europe (Optimer clinical trial) (Dingle et al., 2017; Stoesser et al., 2017; Knetsch et al., 2014; Louie et al., 2011; Cornely et al., 2012). The total number of isolates of each genotype is shown above the bar. **(B) The numbers of genomes in the collections above which contained additional non-*tetM* tetracycline resistance determinants.** For the ST11(078) genotype, the additional Scottish (n=110) isolate collection was also included (indicated by +110 above the bar). Therefore, the total ST11(078) isolates examined was 340 isolates; (the n=230 included in (A) above; plus additional Scottish ST11s (n=110)) the aim being to illustrate the overall prevalence of ‘non-*tetM’* tetracycline resistance determinants within this genotype.

Genotypes could be classified as (i) >60% *tetM* positive, (ii) >0% but <20% *tetM* positive (majority <5%) or (iii) *tetM* not detected (Figure 1A). Non-RT078 genotypes were <20% *tetM* positive, with the notable exceptions of ST37(017) (52/80, 65.0%), ST54(012) (35/58, 60.0%), ST35(046) (27/39, 69.2%), and ST48 (8/12. 67.0%), plus non-toxigenic ST26(140) (65/65, 100%) (Figure 1A). Therefore, at over 75%, RT078 was the most *tetM* positive clinically relevant genotype.

### Prevalence of additional tetracycline resistance determinants in the clinical *C. difficile* population

Each genotype was assessed for the presence of additional tetracycline resistance determinants (Table 1, Figure 1B). The tetracycline efflux pump gene *tet40* (not previously described in *C. difficile*) was present in 27.3% (109/400) RT078 (Figure 1B), and non-*tetM* ribosomal protection proteins were present in 5.5% (22/400) RT078 (Figure 1B). In contrast, only one or two genomes of other ST/PCR-ribotypes were positive for alternative tetracycline resistance determinants, except ST54(012) (n=54), in which eight examples of four additional tetracycline resistance determinants were found and ST33(216), which contained 10/25 (40%) *tetAB(P)* (Figure 1B).

### UK-representative RT078 Phylogeny

A UK-specific RT078 phylogeny was constructed using genomes from clinical infections in Oxfordshire (n=78), Leeds region (n=104) and Scotland (n=110) (Figure 2A, Table S1). Annotation revealed minimal evidence of geographic structure, contrasting markedly with the highly structured distribution of *tetM* sequences (Figure 2B, C) described in detail below.

**Figure 2.**
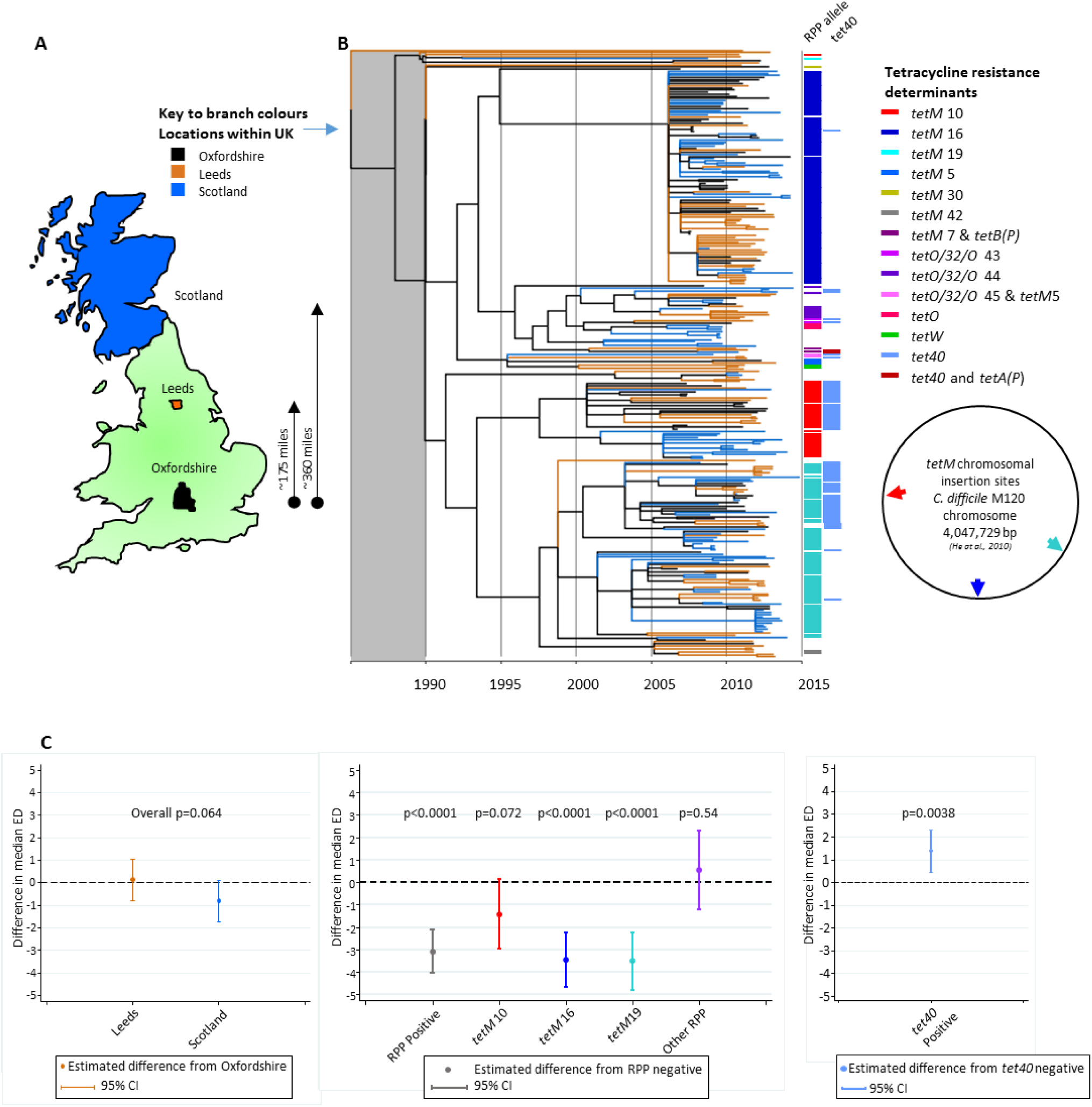
UK-representative, time-scaled RT078 phylogeny revealing a lack of geographic structure, but strong structuring of tetracycline resistance. **(A)** Map showing areas of the UK from which the RT078 *C. difficile* genomes were obtained. **(B)** Time-Scaled ClonalFrameML phylogeny constructed using genomes from UK *C. difficile* isolates; Oxfordshire (n=78), Leeds (n=104), and Scottish (n=110). Branch colours, as per (A), denote the location of each genome. Coloured bars to the right of the phylogeny indicate the presence of tetracycline resistance determinants; ribosomal protection protein allele sequences detected within each genome, were assigned numbers to identify distinct nucleotide sequences of *tetM, tetO/32/O, tetO* or *tetW*. To the right of the phylogeny, the chromosomal location of the three most prevalent *tetM* alleles (designated *tetM* 10, 16 and 19) relative to the RT078 M120 genome (NCBI Reference Sequence NC_017174.1) are shown. All phylogenies included in this study are directly comparable post 1990 ie the timeframe of RT078 emergence; the grey shaded block over the region prior to this date indicates the region which is not scaled identically and should not be compared. **(C)** Extent to which RT078 clonal expansions are associated with geographic structure and tetracycline resistance (ribosomal protection proteins and efflux pumps), using two-sided quantile regression. (i) Difference in median Evolutionary Distinctiveness score compared to Oxfordshire samples. A lower Evolutionary Distinctiveness value indicates a larger proportion of close relatives in the tree. The p-value measures the overall significance of geographic location on Evolutionary Distinctiveness score. (ii) Difference in median Evolutionary Distinctiveness score for samples with ribosomal protection proteins detected compared to ribosomal protection protein-negative samples, overall and for each of the three putative *tetM*-associated clonal expansions. A lower Evolutionary Distinctiveness value indicates a larger proportion of close relatives in the tree. The p-values measure the significance of gene presence on Evolutionary Distinctiveness score. (iii) Difference in median Evolutionary Distinctiveness score for samples with tetracycline efflux pumps (*tet40* and *tetA*(*P*)) detected compared to efflux pump negative samples. A lower Evolutionary Distinctiveness value indicates a larger proportion of close relatives in the tree. The p-value measures the significance of gene presence on Evolutionary Distinctiveness score.

Prior to annotation, distinct *tetM* allele sequences were assigned a number (available at https://pubmlst.org/bigsdb?db=pubmlst_cdifficile_seqdef&page=downloadAlleles>). Among the *tetM* positive UK RT078s, three *tetM* alleles predominated; *tetM* 10 (36/292, 12.3%), *tetM* 16 (101/292, 34.6%) and *tetM* 19 (78/292, 26.7%) (Figure 2B). Coloured bars (or branches Figure S1) were used to identify distinct *tetM* alleles (Figure 2B).

*tetM* alleles 10, 16 and 19 were each carried by closely related, Tn916-like conjugative transposons. Independent acquisition events, estimated from the phylogeny to have occurred between 1995–2006, were suggested by their unique chromosomal insertion sites (Figure 2B). Acquisition of *tetM* 16 or *tetM* 19 was associated with significantly shorter branch lengths (confirmed by median Evolutionary Distinctiveness scores equalling 3.78 and 3.58 respectively, versus 7.22 for branches representing genomes lacking a ribosomal protection protein gene, p<0.001). This observation is consistent with clonal expansion in response to tetracycline-associated selection pressure (Figure 2B, C); significantly lower ED scores indicate unexpectedly short branches (Isaac et al., 2007). It is possible that for a given branch there could be some other genetic change other than *tetM* that is the cause of the clonal expansion, but since we see the same pattern on several independent branches where *tetM* was acquired, it seems very likely that this underlies the clonal expansion. The acquisition of efflux pump *tet40* on its own was not associated with clonal expansion, with only a slightly higher median Evolutionary Distinctiveness score compared with *tet40* absence (Figure 2C).

The same phylogeny was annotated for the presence of additional resistance determinants (conferring aminoglycoside, fluoroquinolone or clindamycin resistance, Figure S2A–C), but no evidence of associated clonal expansions was found independent of the *tetM*-associated expansions (Figure S2D–F).

### RT078 Phylogenies representing UK Regions

Separate phylogenies were constructed to examine the detailed evolutionary history of RT078 within two of the geographic regions represented in the UK-phylogeny. These were Scotland (population 5.295 million, area 30,918 square miles) (Figure 3A, B) and Oxfordshire (population 655,000; area 1,006 square miles) (Figure 3A, C).

**Figure 3.**
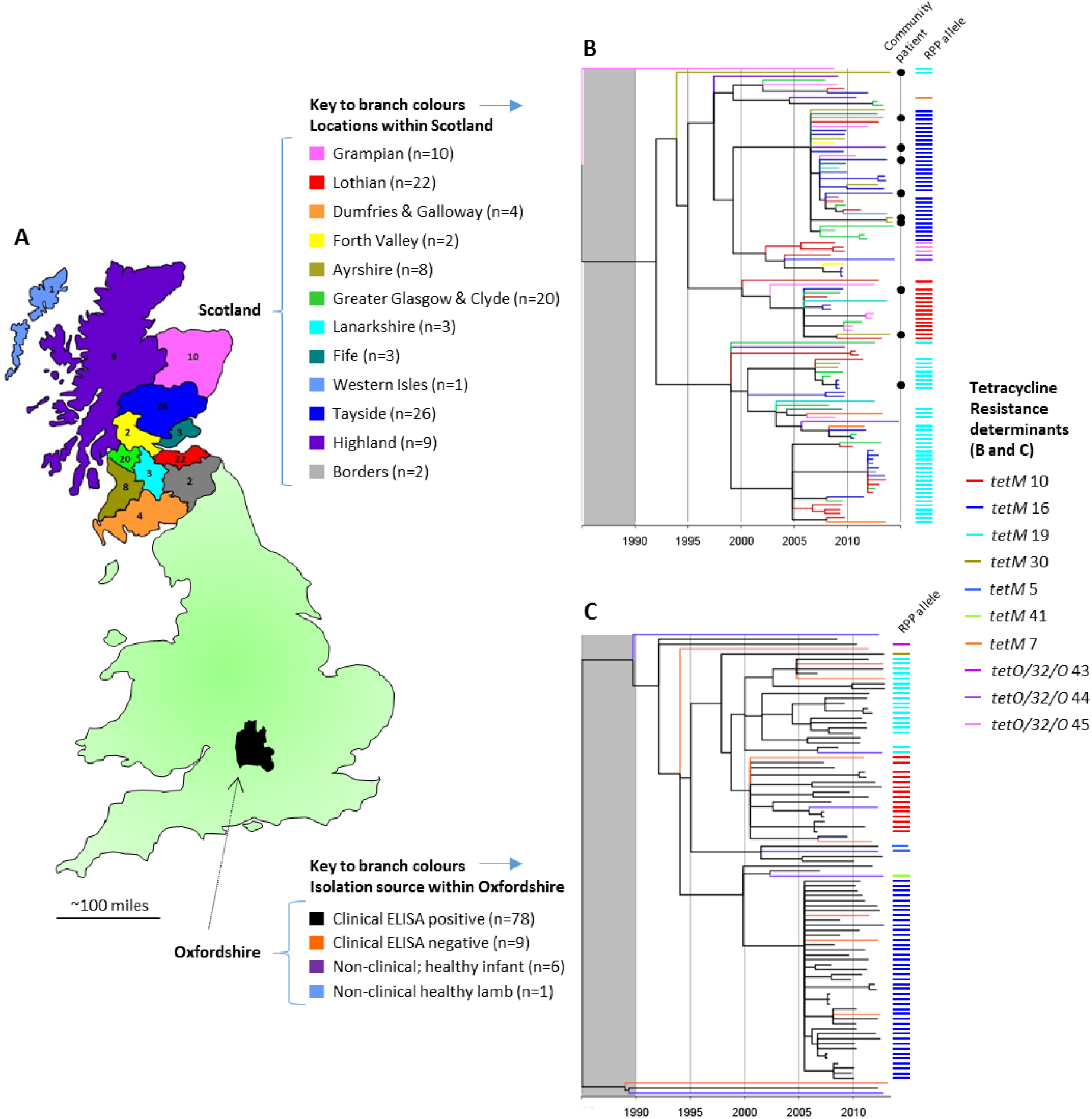
UK Regional RT078 Phylogenies; Scotland and Oxfordshire. (**A**) Map and legend indicating the regions of Scotland and Oxfordshire from which genomes originate. Scottish regions correspond to administrative areas known as ‘health boards’. (**B**) Time-Scaled RT078 phylogeny for Scotland. Branch colours, as per map (A). Coloured bars to the right of the phylogeny denote the ribosomal protection protein (RPP) allele sequences detected within each genome (as Figure 2), numbers being assigned to identify distinct nucleotide sequences of *tetM* or *tetO/32/O*. Isolates were cultured from human clinical samples received from both hospital and community patients, the latter indicated by a black dot. The grey shaded block over the region prior to 1990 indicates the region which is not scaled identically for different phylogenies and should not be compared. (**C**) Time-Scaled RT078 phylogeny for Oxfordshire clinical and non-clinical isolates. Branch colours as per map (A). Coloured bars indicate ribosomal protection protein alleles as above.

The branches of the Scottish phylogeny (n=110 genomes, Figure 3B) were coloured to represent geographic regions (administrative areas, or ‘health boards’, Figure 3A), thus increasing the level of geographic discrimination. As before, geographic structure was absent, healthcare-associated and community isolates intermingling (Figure 3B), but the distribution of the *tetM* alleles 10, 16 and 19 within the phylogeny was highly structured.

The Oxfordshire regional phylogeny (n=94 genomes, Figure 3C), represented a more densely sampled, smaller geographic area (Figure 3A). Here, the EIA-positive *C. difficile* clinical isolate genomes (n=78, as in Figure 2B) were supplemented with EIA-negative clinical isolates (n=9 i.e. isolates patients with diarrhoea but without evidence of toxin production suggesting *C. difficile* was colonising the patient rather than causing disease), non-clinical isolates from healthy infants (n=6) (Stoesser et al., 2017) and a lamb (n=1) (Table S1). All genomes were pathogenicity locus ie. toxin A and B encoding sequence positive (Braun et al., 1996) This regional phylogeny also lacked structure according to location or isolation source, but it was again structured according to *tetM* allele (Figure 3C).

### International phylogenies confirm three *tetM* positive RT078 clades are present across continents

Two international RT078 phylogenies were constructed using genomes from clinical infections in England, (Oxfordshire n=78 and Leeds n=104) supplemented firstly with clinical and non-clinical isolates from The Netherlands (Knetsch et al., 2014) (Table S1, human clinical n=25, human farmers n=15, pig isolates n =20) (Figure 4A, B), and secondly, with genomes from clinical infections in North America (Table S1, USA n=15 and Canada n=4) and Europe (five countries, n=13) (Figure 4C, D). Once again, structure according to geography was absent, but structuring by *tetM* allele (Figure 4A, B) was clear.

**Figure 4.**
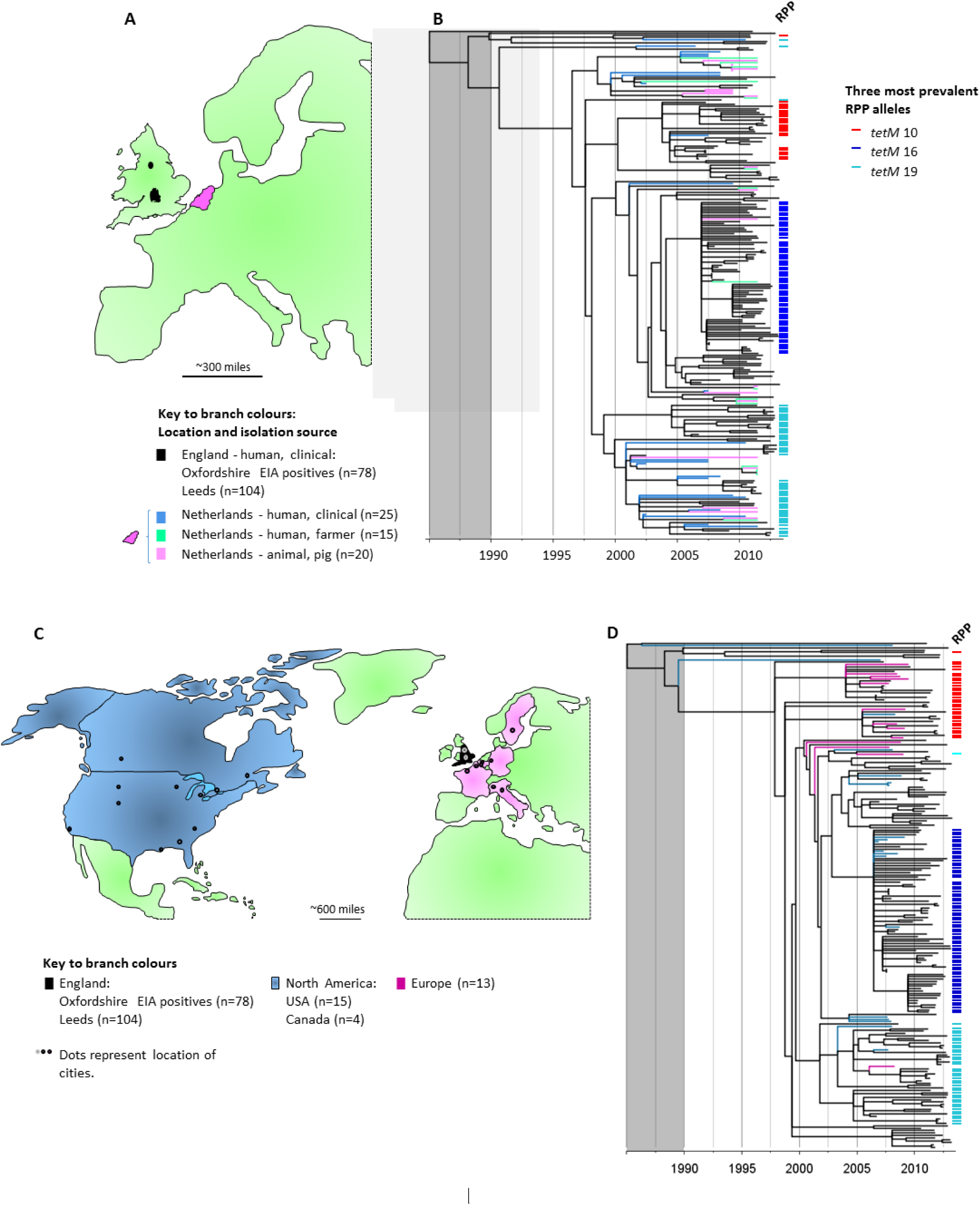
International phylogenies confirm three major *tetM* positive RT078 clades are present across continents. **(A)** Map of Western Europe; regions of England and The Netherlands, from which the genomes included in (B) originate are highlighted (black and pink respectively). (**B**) Time-Scaled RT078 phylogeny constructed using genomes of clinical isolates from England (Oxfordshire and Leeds), supplemented with genomes from the Netherlands (human clinical, farmer and pig isolates (Knetsch et al., 2014). Branch colours as per map (A). The presence of the three predominant ribosomal protection protein (RPP) *tetM* alleles (*tetM* 10, 16 and 19) is indicated by the coloured bars to the right of the tree. The grey shaded block over the region prior to 1990 indicates the region which is not scaled identically for different phylogenies and should not be compared. (**C**) Map highlighting North America and Western Europe, from which the genomes included in (D) originate. (**D1**) Time-Scaled RT078 phylogeny constructed using genomes of clinical isolates from England supplemented with clinical isolates from North America and Europe (distinct from the isolates used above in (B), from two clinical trials of the drug fidaxomicin (Table S1) (Louie et al., 2011; Cornely et al., 2012).

### Tetracycline selection in other *C. difficile* genotypes

Over 60% of genomes belonging to each of five non-RT078 genotypes were *tetM* positive (Figure 1A). Four of these were investigated phylogenetically; ST37(017), ST54(012) and ST35(046) and non-toxigenic ST26(140), (Table S2, Dingle et al., 2017; Stoesser et al., 2017), but not ST48(038/104, as only 12 genomes were available. Branches were coloured according to isolation source and geography as before, and *tetM* alleles indicated by coloured bars (Figure 5A–D). As previously for RT078, additional resistance determinants, (Table 1), were also highlighted (when present in five genomes or more), to reveal the possible impact of selection by other antimicrobials (coloured dots, Figure 5A–D).

**Figure 5.**
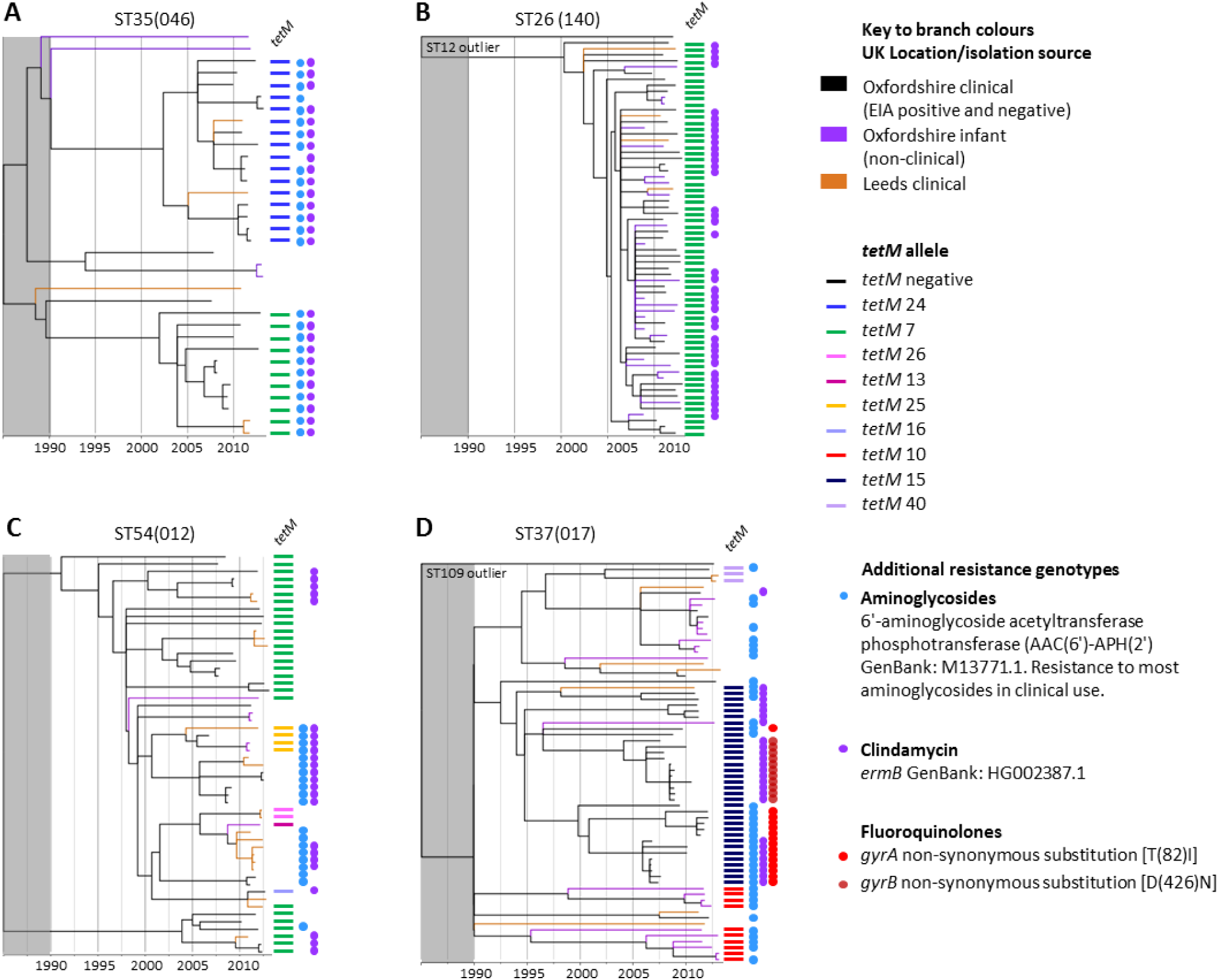
Phylogenetic analysis of additional *tetM* positive *C. difficile* genotypes. Time-scaled phylogenies were constructed representing four non-RT078 genotypes with >60% *tetM* prevalence; **(A)** ST35(046), **(B)** ST26(140), **(C)** ST54(012) and **(D)** ST37(017). In phylogenies (B) and (D) a single closely related genome of a distinct genotype (ST12 and ST109 respectively) was included to ensure that the tree was rooted pre-1990 and that the four phylogenies could therefore be compared post 1990. The grey shaded block over the region prior to 1990 indicates the region which is not scaled identically for different phylogenies and should not be compared. Genomes were from Oxfordshire (clinical EIA positives and negatives plus non-clinical, healthy infants) and Leeds (clinical isolates); branch colours indicate location/isolation source as before. Coloured bars to the right of each phylogeny indicate the presence of tetracycline resistance determinants. Coloured dots represent additional genetic determinants identified, conferring resistance to fluoroquinolones, rifampicin, clindamycin and aminoglycosides (Table 1), which are shown if five or more positive genomes were identified per genotype.

A number of recent *tetM* acquisition events were obvious (Figure 5). These were followed by possible clonal expansions most notably within genotypes ST35(046) and non-toxigenic ST26(140) (Figure 5A, B). Clonal expansion was particularly marked in ST26(140), where all genomes were *tetM* positive, and clonal expansion occurred in the absence of disease-causing ability, this genotype being non-toxigenic (lacking the pathogenicity locus, PaLoc, in all genomes (Braun et al., 1996)). With the exception of ST35(046) where aminoglycoside and clindamycin resistance determinants co-localised with *tetM* (Figure 5A), and the fluoroquinolone resistant region of the ST37(017) phylogeny (Figure 6D, Dingle et al., 2017), there was no clear evidence of clonal expansions which had followed the acquisition of the non-*tetM* antimicrobial resistance determinants. In common with RT078, all four phylogenies (Figure 5) lacked geographic structure, with the exception of the fluoroquinolone resistance region of the ST37(017) phylogeny (Figure 5D) (Dingle et al., 2017).

### Sequences of RT078 tetracycline resistance determinants support its zoonotic origin

The *tetM* sequences described here in *C. difficile* are typical of many Gram-positives, including established zoonotic species. For example, the RT078 *tetM* 10 allele shared 100% nucleotide sequence identity with *tetM* genes of *Streptococcus agalactiae, Enterococcus faecalis* and *Streptococcus pneumoniae*, and 99% nucleotide sequence identity with *Streptococcus suis* (a pathogen of pigs that can infect humans occupationally (Hoe at al., 2011; Mancini et al., 2016)). Identical *tetM* 10 sequences have also been found in Gram-negative bacteria including *Escherichia coli*. The RT078 *tetM* 16 allele shared >97% nucleotide sequence identity with *tetM* in *Enterococcus* species (mostly *E. faecalis* and *E. faecium*), followed by *Staphylococcus* and *Streptococcus* species, including *S. suis*.

Other RT078 tetracycline resistance determinants were also identical, or very closely related to those found in zoonotic bacteria. For example, RT078 *tet40* sequences shared 99–100% identity with *Streptococcus suis tet40* (GenBank KC790465.1) and the RT078 *tetO* sequences shared over 99% nucleotide sequence identity with *Campylobacter jejuni, Campylobacter coli* and *S. suis tetO* sequences. In addition, the RT078 *tetO/32/O* mosaic sequence shared 99% identity with the sequence found in the *S. suis* genome.

## Discussion

Our time-scaled phylogenies revealed parallel *tetM*-associated RT078 clonal expansions, dating from around the year 2000 (Figures 2B, 3B,C, 4B,D). These findings are consistent with an evolutionary response to tetracycline selective pressure, during the timeframe of increasing RT078-associated clinical cases (Goorhuis et al., 2008; Bauer et al., 2011; Burns et al., 2010; Mulvey et al., 2010; Health Protection Scotland, 2017). Tetracyclines were initially introduced around 60 years ago in both clinical and veterinary settings. However, following the emergence of resistance they were largely replaced in human medicine by fluoroquinolones (Thaker et al., 2010). By 2010–13 tetracyclines comprised <18% of total antibiotics consumed in England, and 92% of this was in general practice (Public Health England, 2014). Over the time period relevant to this study, tetracyclines were most commonly used for the treatment of acne and chlamydial sexually transmitted diseases. It is implausible that such prescribing in teenagers and young adults provided extensive selection pressure for *C. difficile*, given that healthy individuals living in the community have very low colonisation rates for these bacteria (Manzoor et al., 2017). This study therefore provides evidence of a plausible agricultural link underlying the emergence of RT078, by presenting its recent evolutionary history with respect to the acquisition of antimicrobial resistance.

In contract to humans, tetracyclines remain the most widely used antimicrobial for the treatment of infections in animals (European Medicines Agency (2016). In addition, their use as growth promoters (in sub-therapeutic doses) continues outside the EU, this being the only region to ban the practice (Scheel 1970). During 2015, 6,880 metric tonnes of tetracyclines were sold in the USA (US Food and Drug Administration 2015) (representing a 31% increase from 2009), compared to 166 tonnes in the UK (Veterinary Medicines Directorate 2015). The extent of agricultural tetracycline use, the prevalence of RT078 in animals used for food (Hensgens et al., 2012; Keel et al., 2007; Curry et al., 2012; Songer et al., 2007), and the timeframe of RT078 emergence all implicate tetracycline use in agriculture as a plausible source of selective pressure. The global food chain provides an obvious route for rapid RT078 dissemination, but indirect transmission from the agricultural environment to humans via contaminated water or vegetables (Xu et al., 2014) is also possible.

The absence of geographic structure within our RT078 phylogenies is consistent with its rapid international spread (Figure 4), as is the absence of large-scale, localised nosocomial outbreaks (Figures 2B, 3B,C, 4B,D). Among other genotypes, such outbreaks have been associated with extensive prescribing of, and resistance to, high risk antimicrobials such as clindamycin, cephalosporins and fluoroquinolones. In our study, large scale clonal expansions were not associated with fluoroquinolone or clindamycin resistance in RT078 (Figure S2). Equivalent analysis for cephalosporins cannot be performed because the genetic mechanism(s) of resistance in *C. difficile* have yet to be defined and although MICs can vary, *C. difficile* has typically been considered inherently cephalosporin resistant, irrespective of genotype (Shuttleworth et al., 1980). The international spread of RT078 indicates that changes in antimicrobial resistance phenotype could potentially impact at any location, depending on local prescribing practices. Consequently, ST11(126), a frequently fluoroquinolone resistant descendant of RT078 (Figure S1), which is prevalent in Italian clinical settings (Spigaglia et al., 2010; Freeman et al., 2015) is of particular concern, as is the epidemic multidrug resistant RT078 observed in Spanish swine (Peláez et al., 2013).

The identification of widespread tetracycline resistance (>60% *tetM* positive) in only five *C. difficile* genotypes besides 078 (Figure 1A) is consistent with previous reports (Barbut et al., 2007; Huang et al., 2009; Noren et al., 2009). Phylogenetic analysis (Figure 5) showed that ST35(046) contained two plausible *tetM*-associated clonal expansions (Figure 5A), but the relatively small numbers precluded quantitative ‘Evolutionary Distinctiveness’ analysis. Like RT078, ST35(046) has been found in pigs and has caused human outbreaks of *C. difficile* infection (Akerlund et al., 2011). This genotype also illustrates the possibility that selection by one antimicrobial can drive the acquisition of further, linked antibiotic resistance genes, as almost every *tetM* positive ST35(046) genome was also positive for clindamycin and aminoglycoside resistance determinants (Figure 5A). Non-toxigenic ST26(140) contained *tetM* in every genome examined, suggesting stable integration predating a recent clonal expansion (Figure 5B), concurrent with that of RT078. ST26(140) therefore illustrates the consequences of tetracycline selection in a harmless commensal organism, confirming that tetracycline selection may have been sufficient to drive the emergence of RT078.

RT078 has many resistance determinants in common with zoonotic pathogens such as *Streptococcus suis, Campylobacter jejuni* and *C. coli*. Quantitative analysis using ‘Evolutionary Distinctiveness’ (Figure 2C, 2SE–F), confirmed that *tetM* was associated with *C. difficile* 078 clonal expansions. Although widespread in RT078, the tetracycline efflux pump *tet40* did not on its own show such as association (Figure 2B,C). Efflux pumps often confer a low-level resistance phenotype, assisting bacterial survival at sub-lethal concentrations of antimicrobials (for example *tetK* in LA-MRSA CC398, Larsen et al., 2016). They thereby function in promoting the acquisition of further high level resistance determinants, such as *tetM*. The parallels between RT078 and zoonotic *Streptococcus suis* also extend to their epidemiology. *S. suis* is a globally distributed emergent pathogen of humans (Werteim et al., 2009), commonly isolated from pigs, geographic clustering of subpopulations is absent (Weinert et al., 2015) and *S. suis* exhibits rapid, recent increases in tetracycline resistance (Hoa et al., 2011). The emergence of human pathogens, coincident with tetracycline resistance acquisition, has also been noted among other bacterial species. In Group B streptococcus, tetracycline resistance may have contributed to its emergence as a leading cause of human neonatal infections (da Cunha et al., 2014). Among livestock-associated methicillin-resistant *Staphylococcus aureus* CC398, almost all isolates are *tetM* positive, and often also carry the *tetK* (efflux pump) (Larsen et al., 2016).

Although multiple lines of evidence indicate a role for tetracycline selection in the recent evolutionary history of RT078, the possibility exists that further genetic changes (unrelated to tetracycline resistance) contributed to its *tetM*-associated clonal expansions (Figures 2–4, Figure S1). The role of selection by other antimicrobials (fluoroquinolones, clindamycin, aminoglycosides) was investigated (Figure S2A–F), and a small potential contribution by aminoglycosides was indicated by the presence of the resistance gene *aphA1* in a minority of RT078 (Figure S2A). However, *aphA1* could not be assessed independent of *tetM* (Figure S1) because the two genes co-localised. Further work would be required to compare the total gene content of *tetM* positive RT078 isolates with older *tetM* negatives, to identify further potentially relevant genetic differences that could explain the clonal expansions. The identification of *tetM*-associated clonal expansions in genetically divergent *C. difficile* genotypes; ST35(046) (together with clindamycin and aminoglycoside resistance determinants, Figure 5A) and non-toxigenic ST26(140) (Figure 5B), serve to further highlight *tetM* as a factor common to recent clonal expansions within distinct *C. difficile* genetic backgrounds. To further confirm the zoonotic origin of RT078, additional data on the changing use of tetracycline over time, and concurrent isolates from clinical cases and farm animals would be useful, although challenging to source retrospectively.

In summary, numerous lines of evidence support the hypothesis that tetracycline use, plausibly in agriculture, has provided recent selection pressure which has impacted on the evolution of tetracycline resistant RT078. The major *C. difficile* RT078 transmission routes to humans are consequently more likely to be related to agriculture and international food chains than nosocomial. Our work strongly suggests that the use of antimicrobials outside the healthcare environment can not only select for resistant organisms, but can contribute to the emergence of human pathogens. Furthermore, our findings add to the body of evidence supporting initiatives such as ‘One Health’ (One Health Initiative, 2018), which aims to expand interdisciplinary collaboration in all aspects of health care for humans, animals and the environment.

## Methods

### *C. difficile* Whole Genome Sequences

*C. difficile* genomes derived from isolates of either RT078 or ST11 (n=400) were sourced from several published collections (Dingle et al., 2017; Stoesser et al., 2017; Knetsch et al., 2014; Louie et al., 2011; Cornely et al., 2012), as well as from an unpublished Scottish collection and the Oxford University farm, Wytham, UK; Table S1 and items (i) to (v) below. Each isolate was obtained from a distinct sample. EIA-negative isolates were inferred to be toxigenic or non-toxigenic, depending on the presence/absence of the toxin encoding pathogenicity locus (PaLoc) (Braun et al., 1996) (Tables S1, S2). Complete collections in (i), (iii) and (iv) below have been described previously (Dingle et al., 2017; Stoesser et al., 2017; Knetsch et al., 2014).

### (i) Clinical *C. difficile*: Oxfordshire and Leeds, UK

Genomes were available for 87 RT078 or ST11 *C. difficile* isolates cultured from symptomatic Oxfordshire patients by the Clinical Microbiology Laboratory, Oxford University Hospitals NHS Trust, Oxford between September 2006 – April 2013 (Dingle et al., 2017). Seventy-eight isolates were derived from toxin immunoassay (EIA)-positive stools (initially the Meridian Premier Toxins A&B Enzyme Immunoassay [Meridian Bioscience Europe, Milan, Italy], until April 2012 and subsequently the TechLab Tox A/B II assay [TechLab Inc, Blacksburg, VA, USA]), and nine from EIA-negative, but glutamate dehydrogenase (GDH)-positive (by the Premier *C. difficile* GDH EIA, Meridian Bioscience Europe, Milan, Italy) (Dingle et al., 2017). Genomes were also available for 104 RT078 *C. difficile* positive stool samples (identified using cytotoxin testing) obtained from routinely examined, diarrhoeal faecal samples at the Leeds Teaching Hospitals NHS Trust, between August 2010 – April 2013 (Table S1).

### (ii) Clinical *C. difficile*: Scotland, UK

Isolates from Scotland, UK, included 109 isolates of RT078, and one closely related RT066 isolate (Table S1). These isolates form part of a collection stored at the Scottish Microbiology Reference Laboratory (Glasgow). Cultures are provided by all twelve Scottish regional NHS Healthcare Boards in the event of a severe/fatal case, a suspected outbreak or a suspected ribotype 027 infection. In addition each Health Board provides a fixed number of samples based on the rates of infection/population. This allows surveillance of prevalent circulating strains to be assessed. RT078 isolates for this study were selected based on the ribotypes from samples referred to the Reference laboratory between November 2007 – October 2014, and aiming to provide the widest temporal and geographical representation. Locally, positive faecal stool samples were identified prior to 2009 using a toxin specific EIA (or cell cytotoxicity), and post 2009, using a two step algorithm requiring GDH detection, followed by toxin assessment. These samples were from patients located in healthcare (n=99) and community settings (n=10) (unassigned n=1) from twelve of fourteen Health Boards (as per the map, there were no relevant samples from two small island Health Boards not shown on the map, Figure 3A).

### (iii) Clinical *C. difficile*: North America and Europe

Thirty-two ST11 genomes from North American (Canada n=4, USA n=15) and European (n=13) *C. difficile* isolates cultured from clinical infections between November 2006 – June 2009 were available from a variety of locations from two clinical trials of fidaxomicin (Table S1, showing city and country (Louie et al., 2011; Cornely et al., 2012)). Previously published RT078 genomes from human clinical cases (n=25, 2002–2011) in The Netherlands were also included (Knetsch et al., 2014) (Table S1).

### (iv) Non-Clinical *C. difficile*

Six ST11 isolates were cultured from the stools of healthy, asymptomatic Oxfordshire infants (Stoesser et al., 2017) between April – October 2012 (Table S1). A single ST11 isolate was isolated from a lamb at Oxford University Farm, Oxfordshire, UK, June 2009. Previously published *C. difficile* RT078 genomes from farmers (n=15, 2009–2011) and pigs (n=20, 2008–2011) in The Netherlands were included (Knetsch et al., 2014) (Table S1).

### (v) PCR-Ribotype Reference Isolates

For additional context, five genomes from PCR-ribotype reference *C. difficile* representing RT078, RT126, RT033, RT045 and RT066 were included, all of which are genetically very closely related, sharing the same multilocus sequence type, ST11 (Table S1, Figure S1).

### (vi) Four additional genotypes

Four additional *C. difficile* genotypes were also analysed phylogenetically to contextualise findings in RT078; these were ST35(046) (n=34), ST54(012) (n=54), ST37(017) (n=64) and non-toxigenic ST26(140) (n=65) (Table S2). These four genotypes underwent detailed study because they had the highest *tetM* prevalence after RT078. The isolates came from Oxfordshire EIA positive and negative, Oxfordshire Infant, and Leeds region isolate collections described above (Dingle et al., 2017; Stoesser et al., 2017). ST26(140) lacks the toxin-encoding pathogenicity locus and is therefore carried asymptomatically, providing a naturally occurring ‘control’ for the impact of antimicrobial selection in the absence of disease.

### Genome Assemblies

*C. difficile* genomes were assembled from short reads generated using Illumina technology (Bentley et al., 2008). Reference-based assemblies were made for genomes belonging to *C. difficile* clade 5 (ie RT078 and close relatives) as described (Eyre et al., 2013), by mapping reads to the *C. difficile* M120 reference genome (He et al., 2010) and for non-clade 5 genomes by mapping to the CD630 reference genome (GenBank AM180355.1) (He et al., 2010) (clades as defined, Dingle et al., 2011).

*De novo* assembly was performed using Velvet (version 1.0.7–1.0.18) (Zerbino and Birney., 2008) and VelvetOptimiser 2.1.7 (Gladman and Seeman, 2008), optimising kmer size (k), expected coverage (average kmer coverage of contigs) and coverage cutoff (kmer coverage threshold) to achieve the highest assembly N50 value (length of the smallest contig such that all contigs of that length or less form half of the final assembly).

Reads for unassembled genomes have been submitted to NCBI, BioProject ID number PRJNA304087 (Dingle et al., 2017) and PRJNA381384 for the Scottish isolates (accession numbers provided in Tables S1 and S2).

### Identification of Antimicrobial Resistance Determinants

The *de novo* assemblies were queried using the BLAST function of BIGSdb (Jolley and Maiden 2010) to determine whether genes or non-synonymous point mutations known to confer resistance to antimicrobials including fluoroquinolones, tetracyclines, clindamycin and aminoglycosides were present, and to extract the sequences of interest for further analysis. A list of the resistance gene sequences (and GenBank accession numbers) used to perform the BLAST search is provided (Table 1). For acquired resistance genes, a minimum level of 90% nucleotide sequence identity and gene coverage was required. Each unique *tetM* allele was assigned a number (allele nucleotide sequences available at https://pubmlst.org/bigsdb?db=pubmlst_cdifficile_seqdef&page=downloadAlleles).

### Phylogenetic Analyses

Phylogenetic trees were built, based on the *C. difficile* ST11 reference M120-mapped assemblies, using the maximum likelihood approach implemented in PhyML version 3.1.17 (using a generalized time-reversible substitution model and the “BEST” tree topology search algorithm) (Guindon et al., 2010). The trees were then corrected to account for recombination events using ClonalFrameML (Didelot and Wilson 2015) version 1.11 (using default settings). The nodes of the trees were dated using the previously estimated *C. difficile* evolutionary rate of 1.1 mutation per year (Knetsch et al., 2014) for clade 5 STs (including RT078), and 1.4 mutation per year for all other genotypes (Didelot et al., 2012). The main period of particular interest from 1990 to 2015 was allocated most horizontal space in graphical tree representations by compressing the period pre-1990, making trees directly comparable post-1990. Events before 1990 are not shown since dating older nodes using a short-term evolutionary rate is problematic due to the time-dependency of evolutionary rates (Biek et al., 2015). Graphical representations of trees were made using FigTree version 1.4.2 (Rambaut 2016).

A quantitative assessment of clonal expansion(s) within a phylogeny was performed as described (Isaac et al., 2007) and implemented (Dingle et al., 2017) previously. The Evolutionary Distinctiveness (ED) score of each isolate was calculated, equal to the sum, for all branches on the path from the root to the leaf (isolate), of the length of the branch divided by the number of leaves it supports (Isaac et al., 2007). For a given isolate, a low ED score indicated the presence of close relatives in the tree, whereas a high ED score indicated their relative absence. ED scores were compared across various factors using quantile regression statistics, performed using Stata version 14.1 (College Station, TX, USA).

## Acknowledgements

This project was funded by the National Institute for Health Research Health Protection Research Unit (NIHR HPRU) in Healthcare Associated Infections and Antimicrobial Resistance at Oxford University in partnership with Public Health England (PHE). The views expressed are those of the author(s) and not necessarily those of the NHS, the NIHR, the Department of Health or Public Health England.

This study was also supported by the UK Clinical Research Collaboration (Wellcome Trust [grant 087646/Z/08/Z], Medical Research Council, National Institute for Health Research [NIHR grant G0800778]); NIHR Oxford Biomedical Research Centre, NIHR Oxford Health Protection Research Units on Healthcare Associated Infection and Antimicrobial Resistance (grant HPRU-2012–10041) and on Modelling Methodology (grant HPRU-2012–10080), and the Health Innovation Challenge Fund (a parallel funding partnership between the Wellcome Trust [grant WT098615/Z/12/Z] and the Department of Health [grants WT098615 and HICF-T5-358]).

The identification, isolation and analysis of the Scottish RT078 isolates was funded by a Scottish Infection Research Network (SIRN) grant (ref SIRN/09), awarded to GD and CAM (PIs), JC, DB and UI (co-applicants).

KED was supported by NIHR Oxford Biomedical Research Centre, DWC and TEAP are NIHR senior investigators.

SB was supported as a Daphne Jackson Fellow funded by Medical Research Scotland. UZI was supported by a NERC independent research fellowship (NE/LO11956/1).

The authors thank the Food Animal Initiative in Oxford for access to animals to sample for *C. difficile* at the Oxford University Farm, Wytham.

The authors acknowledge the contribution of the Modernising Medical Microbiology Informatics Group comprising: Carlos Del Ojo Elias, Charles Crichton, Vasiliki Kostiou, and Adam Giess (Nuffield Department of Clinical Medicine, University of Oxford, UK), Jim Davies, (Department of Computer Science, University of Oxford, UK).

The funders had no role in the writing of the manuscript or the decision to submit it for publication.

## Author Contributions

KED, XD, TEAP, ASW, and DWC conceived and designed the study; KED, DWE, NS, CAM, JC, DB, SB, UZI, CG, GD, WNF, MHW, TEAP, ASW, and DWC contributed isolates and DNA sequence data; XD and DWE Performed phylogenetic analysis; XD and DWE provided bioinformatics support; TPQ and ASW Performed Evolutionary Distinctiveness analysis; KED analysed DNA sequence data for antimicrobial resistance determinants; KED, XD, DWE, MHW, TEAP, ASW, DWC wrote the manuscript.

---

**Figure S1.**
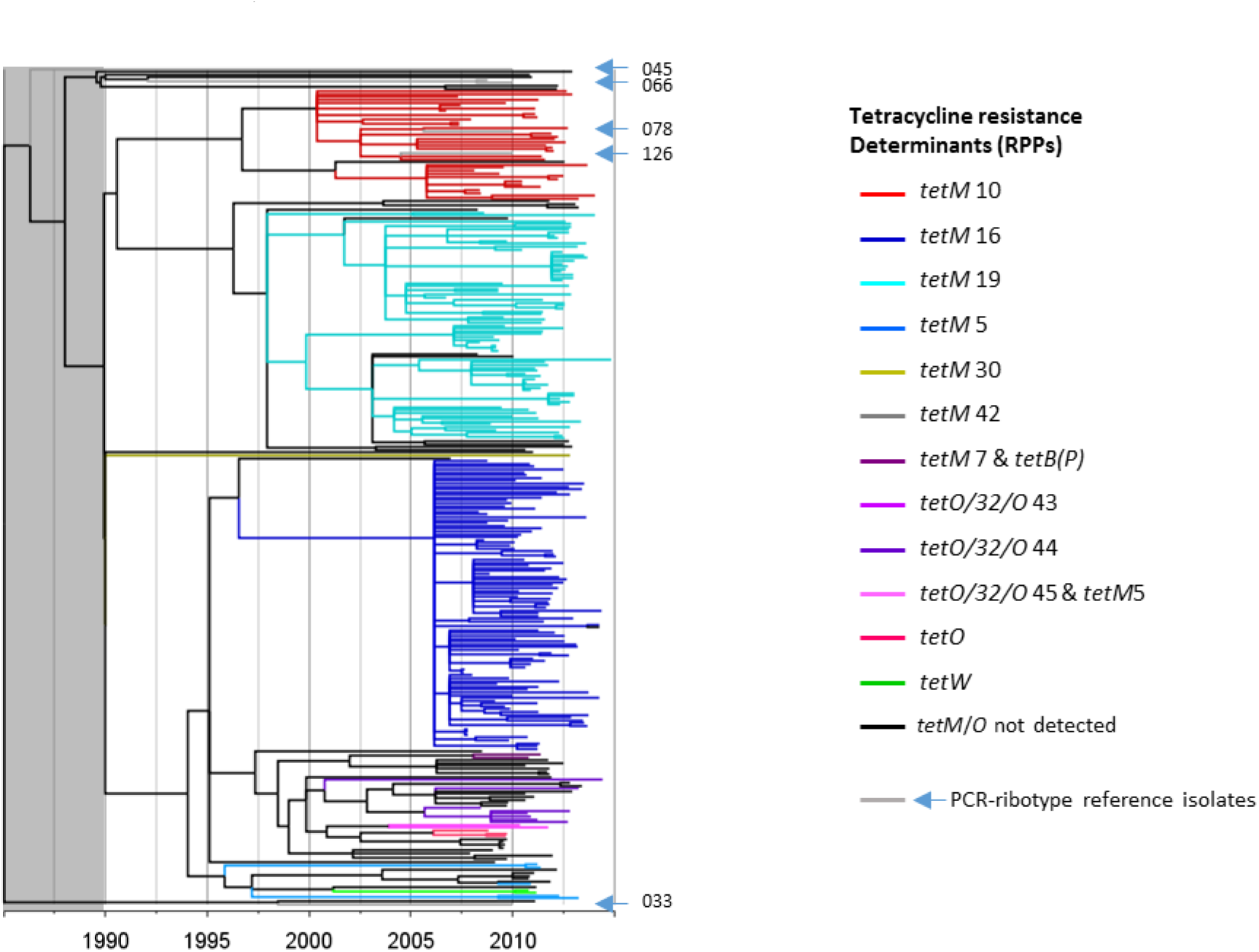
UK-representative, time-scaled ST11/RT078 phylogeny. The phylogeny was constructed using the same genomes as Figure 1, but with an additional five genomes representing five closely related PCR-ribotype reference isolates, all of which are ST11. In contrast to Figure 1 where the branches were coloured for geographic location, here the branch colours indicate presence of tetracycline resistance determinants to highlight the associated clonal expansions.

**Figure S2.**
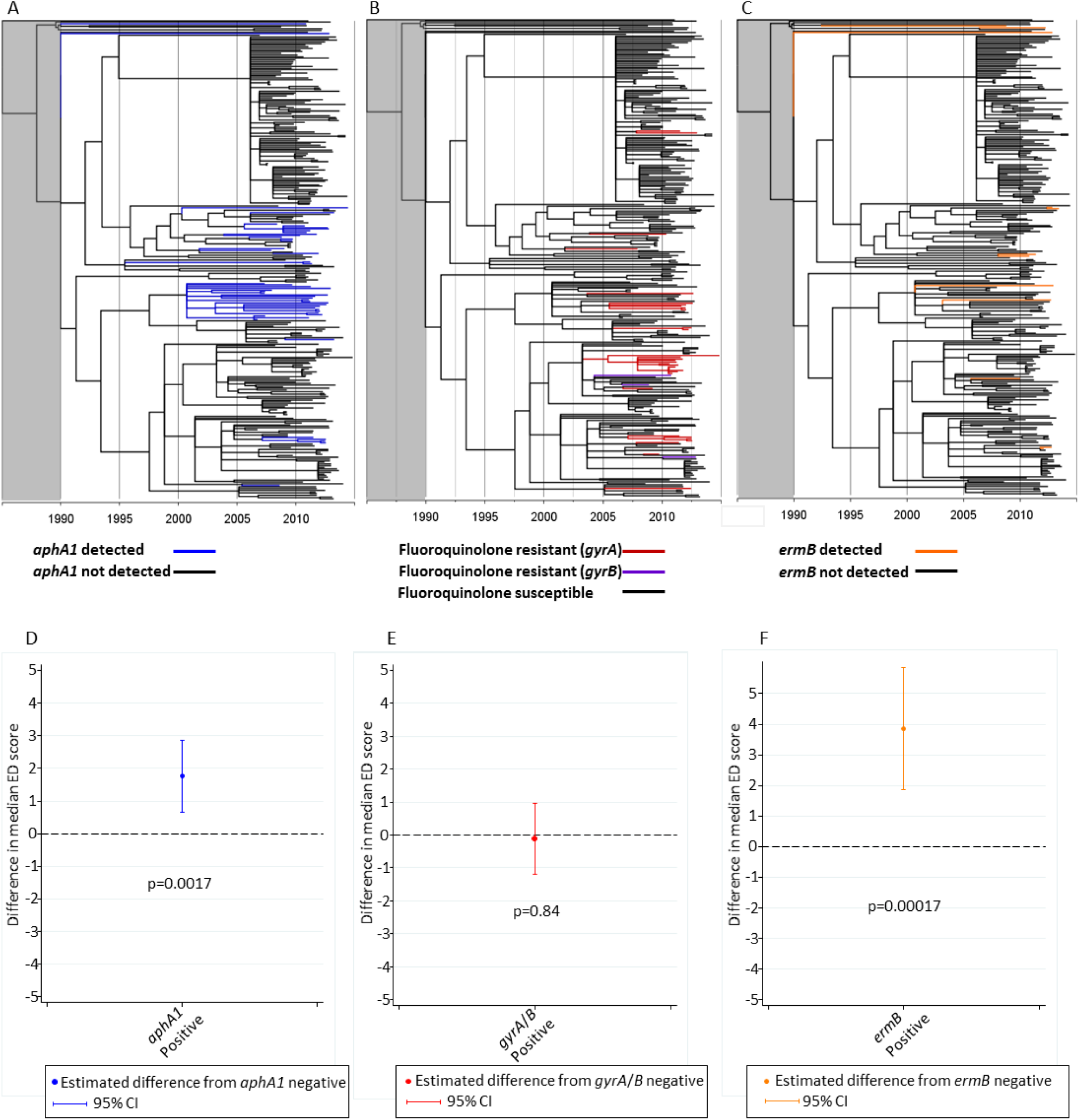
UK-representative, time-scaled RT078 phylogeny. The same phylogeny as Figure 1, however here the branch colours indicate presence of the following antimicrobial resistance determinants: (**A**) Aminoglycoside resistance; *aphA1* (aminoglycoside 3’-phosphotransferase) associated with resistance to streptomycin28. (**B**) Fluoroquinolone resistance conferred by non-synonymous *gyrA* or *gyrB* substitutions, (as cited14). (**C**) Clindamycin resistance; *ermB*. (**D**) Extent to which 078 clonal expansions are associated with antimicrobial resistance determinants *aphA1, gyrA/B* substitutions, and *ermB*, using two-sided quantile regression. Difference in median ED (Evolutionary Distinctiveness) score (Isaac et al., 2007) for isolates with the resistance determinant compared to isolates without. A lower ED value indicates a larger proportion of close relatives in the tree. The p-value measures the significance of the presence of the resistance determinant on ED score.

